# The IMEx Coronavirus interactome: an evolving map of Coronaviridae-Host molecular interactions

**DOI:** 10.1101/2020.06.16.153817

**Authors:** L Perfetto, C Pastrello, N Del-Toro, M Duesbury, M Iannuccelli, M Kotlyar, L Licata, B Meldal, K Panneerselvam, S Panni, N Rahimzadeh, S Ricard-Blum, L Salwinski, A Shrivastava, G Cesareni, M Pellegrini, S Orchard, I Jurisica, HH Hermjakob, P Porras

## Abstract

The current Coronavirus Disease 2019 (COVID-19) pandemic, caused by the Severe Acute Respiratory Syndrome Coronavirus 2 (SARS-CoV-2), has spurred a wave of research of nearly unprecedented scale. Among the different strategies that are being used to understand the disease and develop effective treatments, the study of physical molecular interactions enables studying fine-grained resolution of the mechanisms behind the virus biology and the human organism response. Here we present a curated dataset of physical molecular interactions, manually extracted by IMEx Consortium curators focused on proteins from SARS-CoV-2, SARS-CoV-1 and other members of the *Coronaviridae* family. Currently, the dataset comprises over 2,200 binarized interactions extracted from 86 publications. The dataset can be accessed in the standard formats recommended by the Proteomics Standards Initiative (HUPO-PSI) at the IntAct database website (www.ebi.ac.uk/intact), and will be continuously updated as research on COVID-19 progresses.

## Introduction

Severe acute respiratory syndrome, or SARS, emerged as a life-threatening viral disease of unknown origin in late 2002 in the Guangdong Province of southern China caused by the SARS-CoV-1 virus (1). The Severe Acute Respiratory Syndrome Coronavirus 2 (SARS-CoV-2) is a related virus responsible for the current outbreak of Coronavirus Disease 2019 (COVID-19) (2). As of June 2020, over 7.6 million people globally have been shown to be infected with the virus, with more than 426,000 deaths directly attributed to its effects (John Hopkins University, https://coronavirus.jhu.edu/map.html).

The COVID-19 pandemic has resulted in massive scientific efforts attempting to fight the disease and understand the biology of the virus. This has derived in enormous challenges for the research scientist when attempting to find and select information relevant to specific areas of viral biology and pathology. In order to aid the scientific community and expedite drug and vaccine development, multiple data curation efforts have been undertaken to perform a critical assessment of the literature, and represent different aspects of the virus and the disease in a structured and computationally accessible manner. One recent example of such efforts is the COVID-19 Disease Map (3), a community effort to capture the intricate aspects of SARS-CoV-2 biology as reusable and interoperable pathway maps, so they can be used in systems biology and modelling pipelines.

Identification of virus-host interactions, and the analysis of the topological structure of a relevant molecular interaction network is a necessary step in enabling an understanding of the cellular mechanisms involved in a biological process such as viral infection of a cell. A detailed map of the interactions between human and pathogen proteins will aid a more complete awareness of the mechanisms of infection and subsequent viral replication, assembly and release, and may help to identify novel drug targets, or assist in rapid and more accurate repurposing of existing drugs for treating or preventing infection. Further to this, such networks can be used to study data such as changes in the transcriptome or proteome of a virally infected cell when compared to normal. Co-regulated genes or proteins which also co-cluster in this network may indicate that these entities are involved in the same biological process or are members of the same functional complex.

For such a network to be of value to the researcher, a certain amount of meta-data needs to be provided, which enables the assessment of network quality and data types. These data need to be supplied in a standardized and computer-accessible format, which allows for scoring, filtering and selection. The IMEx Consortium (4) has been providing such data for over 15 years, supplying experimental details using controlled vocabulary terms, captured using a detailed curation model. All aspects of an interaction experiment are described, including host organism, interaction detection and participant identification methodologies and full details of the constructs, such as binding domains and the effects of site-directed mutations. Current membership of the IMEx Consortium includes the IntAct (5), MINT (6), DIP (7), UniProt (8), MatrixDB (9), and IID (10) data resources, who collaborate to provide the users with a single, consistent viral-host dataset to work with. When a novel virus emerges, as was the case with SARS-CoV-2 in 2019, the study of closely related species,such as SARS-CoV-1 and other coronaviruses, may help with this process, giving the scientific community time to produce species-relevant data. For this reason the network includes data on all coronaviruses available in the scientific literature. Whilst primarily consisting of protein-protein interaction data, the network also contains interactions with lipids, glycosaminoglycans and RNAs, again curated to IMEx standards. The data is fully open under a CC-BY 4.0 license and downloadable from the IntAct website and FTP in PSI-MI standard tab-delimited and XML-based formats. A brief description can be found under www.ebi.ac.uk/intact/resources/datasets and a collection of interactive network representations of the dataset at the time of writing is available at http://www.ndexbio.org/#/networkset/4c2268a1-a0f0-11ea-aaef-0ac135e8bacf. The dataset will be expanded and updated with every IntAct release.

## Results and discussion

As of June 2020, the dataset contains 1,778 unique interacting molecule pairs, represented in 2,212 binarized interactions extracted from 86 publications, 5 of which are pre-prints from bioRxiv. The dataset can be downloaded from the IntAct FTP site in PSI-MI standard XML-based formats PSI-MI XML 2.5 (ftp://ftp.ebi.ac.uk/pub/databases/intact/current/psi25/datasets/Coronavirus.zip) and 3.0 (ftp://ftp.ebi.ac.uk/pub/databases/intact/current/psi30/datasets/Coronavirus.zip). It can also be browsed on the IntAct webpage (www.ebi.ac.uk/intact/query/annot:“dataset:coronavirus”), where it is available for download in additional formats, such as the tab-delimited PSI-MI-TAB 2.7. A brief dataset description can be found under www.ebi.ac.uk/intact/resources/datasets#coronavirus. The data can be searched in the IMEx PSICQUIC (11) service and on both the IMEx Consortium webpages (www.imexconsortium.org) and via the VirusMentha (12) browser at https://virusmentha.uniroma2.it/. Interactive network representations for unique molecule pairs and full evidence details can be found in https://doi.org/10.18119/N9MP4S and in https://doi.org/10.18119/N9RC8F. As with all data hosted in IntAct, the dataset is freely available under a CC BY 4.0 license.

The data refers mostly to PPIs (1,674 interactions), plus some interactions involving different RNAs (67 interactions) or small molecules (37 interactions). While data on 70 organisms are included (suppl. Table 1A), most interactions refer to SARS-CoV-2 and SARS-CoV-1 - human interactions (992 and 351 unique interactions, respectively).

IMEx Consortium curators have collated interaction evidence from scientific articles and pre-prints using the following selection criteria:

1. The publication contains interactions involving proteins from any virus member of the *Coronaviridae* family (NCBI taxon ID 11118). This includes not only SARS-CoV-1 and SARS-CoV-2, but also MERS-CoV and members of the family that infect other mammals.
2. The publication contains interactions of human proteins with established relevance for SARS-CoV-2 life cycle (e.g. ACE2 interactions have been included in the dataset).
3. Every interaction described in these publications is curated and included in the dataset, even if their relevance to COVID-19 might seem limited. This results in the inclusion of data from apparently irrelevant species (e.g. yeast) that is of interest from the phylogenetic and evolutionary point of view.
4. Pre-prints are considered and, if deemed appropriate, curated when containing SARS-CoV-2 data, an exception to IMEx practice of representing only peer-reviewed research. This reflects the interest these data have engendered during the pandemic. These datasets are clearly marked as pre-publication and will be re-curated, if necessary, when published.

The IMEx curation model captures the details of specific constructs used for the detection of interactions. This allows for the representation of different construct-associated features, such as specific mutations affecting interaction outcome (13) or sequence regions that are associated with binding. Most of these in-depth studies are centered around the Spike-ACE2 interaction for SARS-CoV-2 and SARS-CoV-1. The only other SARS-CoV-2 mutation reported so far is Nsp5 / 3C-like proteinase p.Cys145Ala catalytically inactive mutant, which exhibits an interaction profile different from the canonical form in an AP-MS study (14).There are a few other mutations reported for SARS-CoV-1 and in other members of the *Coronaviridae* family, but it is clear that specific variation effects on Coronavirus-related interactions is an area requiring extensive exploration. As shown in figures 1B and 1C, fragment constructs have been used in more studies than mutations, although some of these might have been designed for convenience reasons (e.g. constructs that are easier to express in heterologous systems). Interactive network representations highlighting mutations and binding regions can be found in https://doi.org/10.18119/N9W590 and in https://doi.org/10.18119/N90W3W.

**Figure 1:**
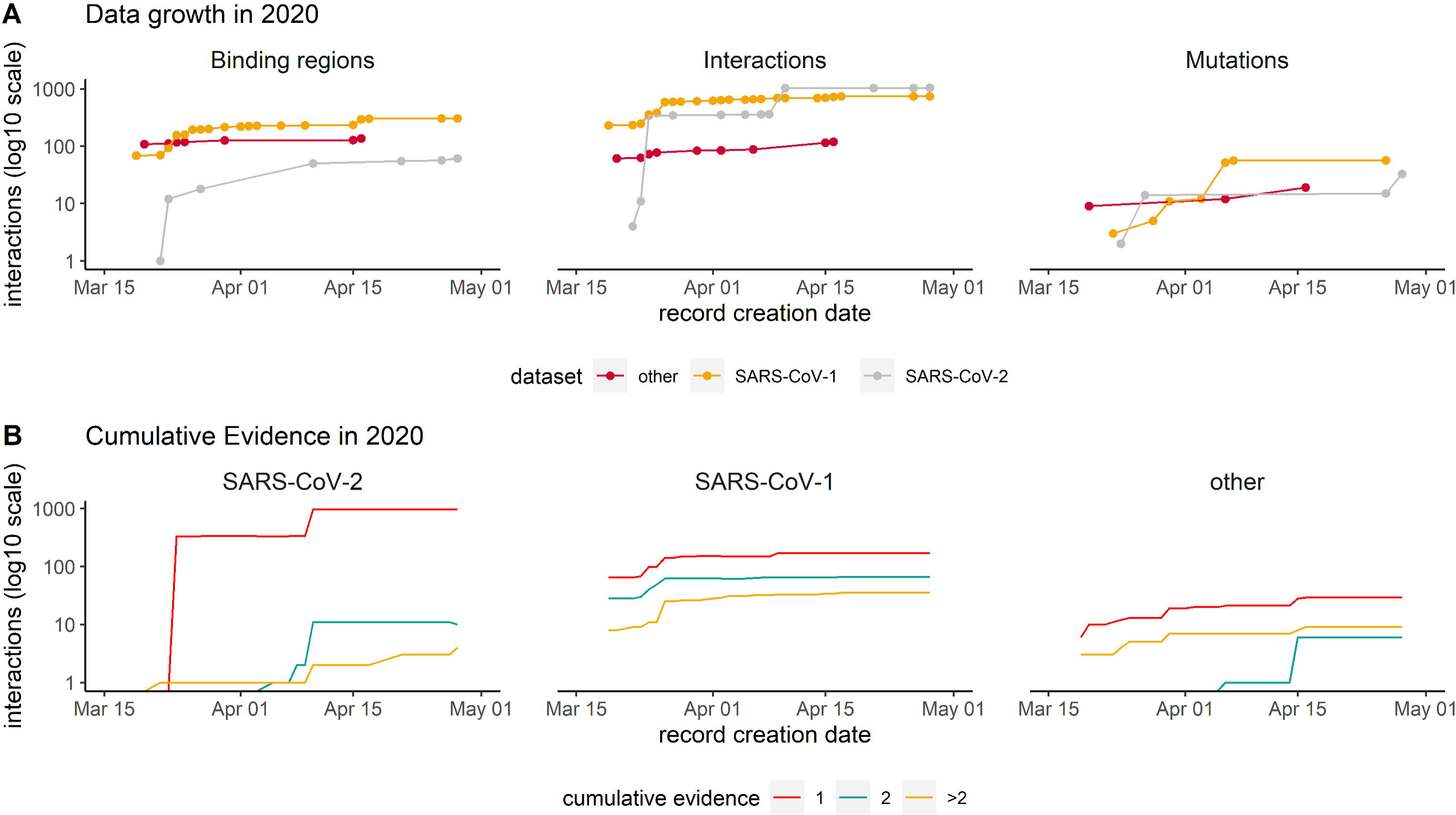
Timeline showing data captured in IMEx resources since COVID-19 outbreak (March 2020). (A) Cumulative interactions, mutation features and binding regions annotated for SARS-CoV-2 (red), SARS-CoV1 (orange) and other *Coronaviridae* family members proteins (grey). Interactions include spoke-expanded binary relationships. Dots represent the date when the interaction was curated. (B) Amount of cumulative experimental evidence associated with unique binary pairs, captured over time, in each of the three dataset: SARS-CoV-1, SARS-CoV-2 and other *Coronaviridae* family members. Interactions include spoke-expanded binary relationships.

Regarding interaction data generation approaches, the experimental setup used to detect an interaction is summarized by three key fields in the IMEx curation model: the method used to determine that an interaction is happening (*interaction detection method*), the method used to determine which molecules are involved (*participant detection method*), and the biological environment in which the interaction occurs (*host organism*). Each of these is described by an appropriate controlled vocabulary term, making the data readily searchable. In the Coronavirus dataset, the overwhelming majority of data were generated by affinity-purification combined with mass-spectrometry approaches performed in HEK293(T) or HeLa cells (suppl. Table 1B). This type of data produces sets of potential interacting partners (preys) associated with a bait of interest, but does not directly identify binary interacting partners. Also, the data needs to be automatically expanded into binaries for its representation in tabular or network graph form. This accounts for a large proportion of spoke-expanded binary relationships found in the dataset: 1,310 out of 2,212 interactions (59%) are expanded binaries, while only 43% of the binary interactions found in the full IntAct database are expanded binaries.

Most content has been curated after declaration of the COVID-19 pandemic on 11/03/2020 (figure 1). The data growth timeline shows three jumps in interaction numbers, two for SARS-CoV-2 and one for SARS-CoV-1, due to curation of high-throughput (HT) studies not available for other *Coronaviridae* (figure 1A). This is also reflected in the pattern of data growth regarding detailed information about mutations and binding regions (figure 1A). The plots also illustrate that at this point there are more studies dealing with detailed interaction data for SARS-CoV-1, a situation that will likely change as more SARS-CoV-2 studies are performed.

Interaction data is derived from both, small-scale studies focused on one or just a few interactions (low-throughput, LT) and large-scale screenings able to detect hundreds or thousands of interactions in a single experiment (HT). The extreme urgency in the study of SARS-CoV-2 has resulted in a strong dominance of high-throughput data for this species in comparison with the other members of *Coronaviridae* reported in the dataset (figure 2A). Additionally, SARS-CoV-2 small-scale studies are mainly focused around the ACE2-Spike interaction due to its relevance for virion recognition and infection. For the remaining interactions, most SARS-CoV-2 data comes from two studies: Gordon et al. (14) and Li et al. (15), both focused on affinity purification techniques combined with mass-spectrometry detection of interacting candidates (AP-MS) in HEK293(T) cells transfected with SARS-CoV-2 proteins. The studies are distinct, showing virtually no overlap (figures 2B-D) and different degree distribution patterns (figure 2E). Gordon et al. shows an unexpected pattern of fully isolated components, likely due to stringent target selection.

**Figure 2:**
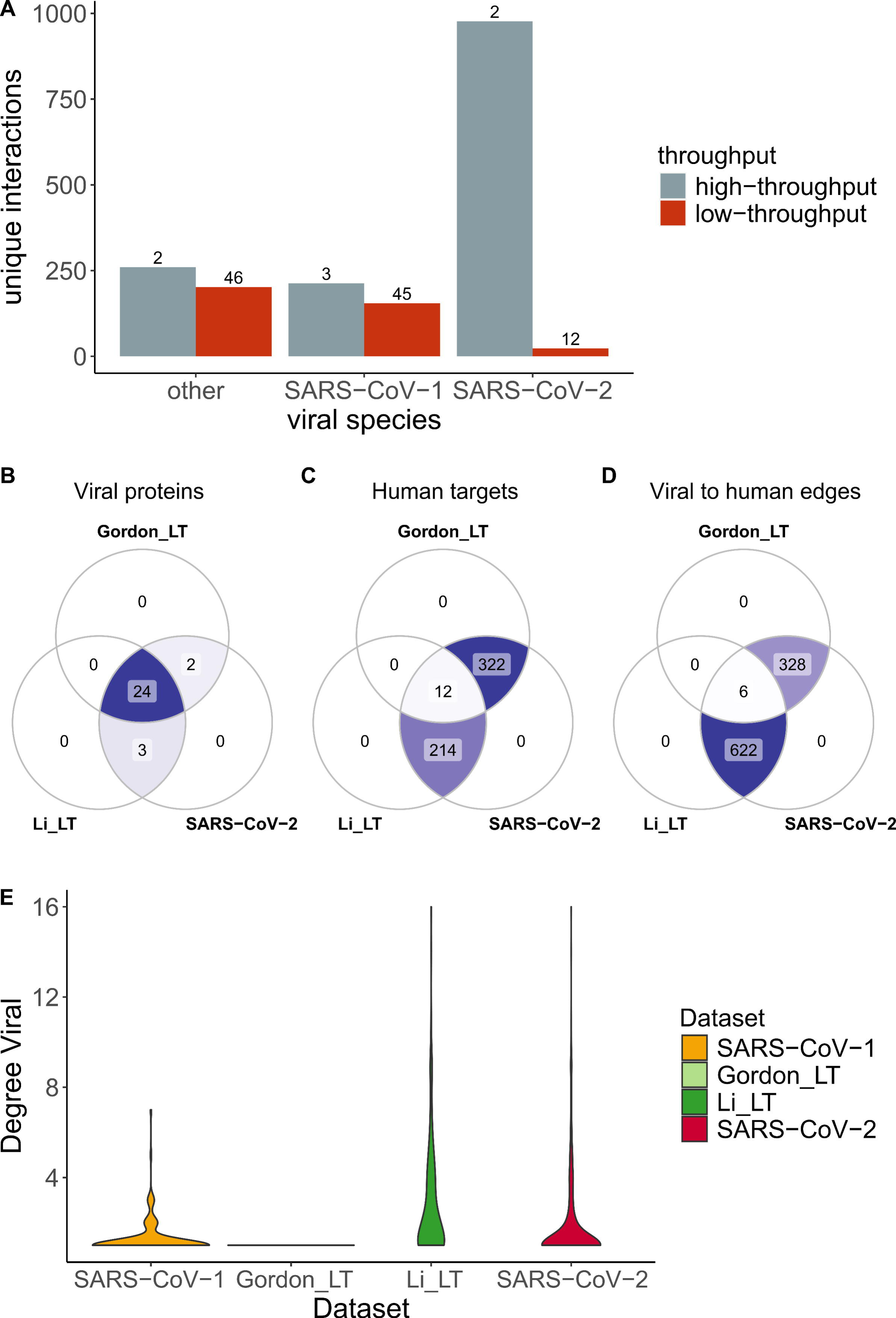
SARS-CoV-2 HT datasets comparison. SARS-CoV-2 datasets include Gordon_LT (Gordon plus low throughput) and Li_LT (Li plus low throughput). (A) Distribution of HT/LT-derived interactions in the Coronavirus dataset. The number on top of each bar indicates the number of publications per category. High-throughput publications are defined as those hosting more than 50 unique interacting pairs. (B-D) Overlap of viral proteins (B), human targets (C) and viral to human edges (D) across datasets. (E) Distribution of number of interactions between viral proteins and human targets. In the y-scale, 1 indicates that one human protein interacts only with one viral protein, while 16 indicates that one human protein interacts with 16 viral proteins.

Lack of overlap between different high-throughput interaction datasets is a long-recognized phenomenon, since different experimental approaches are better suited to detect interactions featuring specific physico-chemical characteristics and protein abundances, among other parameters (16–18). Even in this case, where the approach used is broadly similar, they likely reflect methodological differences. AP-MS datasets are very sensitive to protein abundances and affinities, as well as to the selection and orientation of tags and expression systems. Additionally, strong differences can arise during the selection of *bona fide* interactors, where multiple strategies can be used in order to clean up spurious detections. The lack of common interactions between the different studies focused on SARS-CoV-2 suggest that more of these systematic experiments are needed to increase reliability of biological conclusions extracted from this type of data.

Exploring the biological context of SARS-CoV-2 interactors suggests that the two HT studies are complementary. Pathway enrichment (figure 3 and supplementary table 2) finds commonalities on expected pathways related with cell cycle, response to stress and infectious disease, and DNA and RNA synthesis and processing. Key pathways such as innate immune response or cytokine signaling are only found in one HT study, along with several intracellular signaling and metabolic routes. Only SARS-CoV-1 shows appreciable enrichment in lung tissue, while none of the SARS-CoV-2 sets reach the 5x enrichment threshold suggested by Jain & Tuteja (19) (supplementary figure 1, supplementary table 3). Finally, SARS-CoV-2 interactors from both studies are found as components of histone deacetylase, exosome and ATP-ase transmembrane complexes (supplementary table 4).Including those represented in only one of the studies, we see an abundance of complexes involved in endosome and exosome generation, mitochondrial metabolism, protein production, Ca^2+^-dependent cell signaling and cell cycle control.

**Figure 3:**
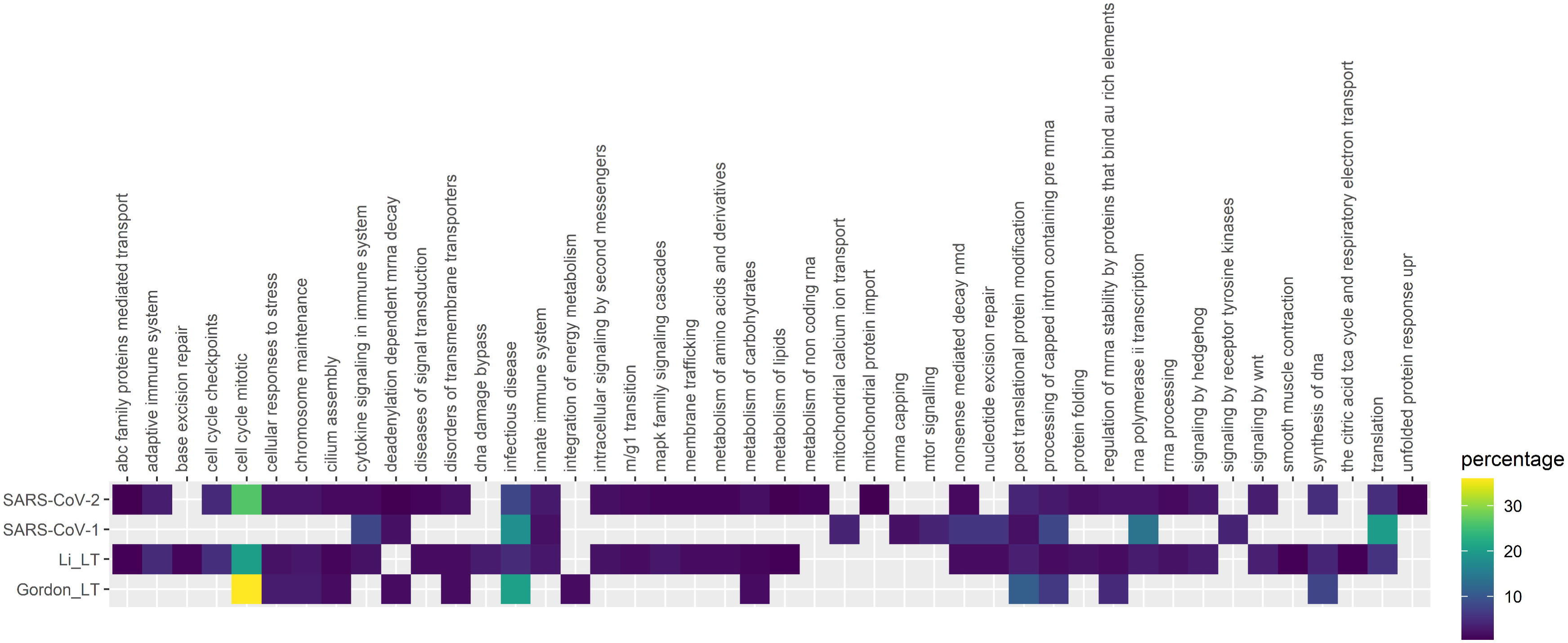
Pathway enrichment analysis of SARS-CoV-1, SARS-CoV-2 and Gordon_LT (Gordon plus low throughput) and Li_LT (Li plus low throughput) datasets. Enrichment was performed using pathDIP (a database integrating 24 different pathway databases). Only human proteins were considered. The majority of enriched pathways were from Reactome database, so a mapping of each Reactome pathway to the parent pathway ontology was performed, and the heatmap shows the percentage of pathways in each parent pathway over the total of pathways.

Development of the IMEx Coronavirus dataset complements other curation efforts that have been initiated in the light of the pandemic. The dataset has been used by the DisGeNET database (20) for contextual annotation with related diseases (https://www.disgenet.org/downloads#section9). Also, members of the IMEx Consortium are involved in the COVID-19 Disease Map initiative (https://covid.pages.uni.lu/), where interaction information from the dataset is guiding COVID-19-related pathway curation. As an example, the list of PMIDs from IMEx Coronavirus dataset has been used to screen papers containing causal interactions to build COVID-19 causal network perturbed during SARS-COV2 infection by the SIGNOR 2.0 resource (21) (https://signor.uniroma2.it/covid/) and to select the Gordon et al. human interactors to integrate in the SIGNOR 2.0 network.

We are also involved in a parallel effort curating SARS-CoV-1 and SARS-CoV-2 related protein complexes in the Complex Portal (www.ebi.ac.uk/complexportal), linking available experimental interaction evidence when possible. This initiative is especially relevant because, as previously stated, coronaviruses increase the number of functional proteins produced by the viral genome by post-translational cleavage of long polypeptide transcripts. The functionality and/or stability of these proteins is further increased through the formation of protein complexes, all of which have been catalogued into the Complex Portal. To date 12 complexes have been identified in each strain, formed by viral-viral protein interactions, including homomeric assemblies such as the dimeric SARS-CoV-2 main protease complex (CPX-5685). Other reference entities such as the SARS-CoV-2 Spike - human ACE2 receptor complex (CPX-5683) have also been created to enable their identification in large - omics datasets. All complexes have been annotated with Gene Ontology terms, describing their role in the virus lifecycle, which again will assist in the analysis of large Omics-derived datasets. Ongoing work includes the addition of the MERS virus complexosome to the available data.

## Conclusions

Accurate and detailed representation of biological insight into public databases is a fundamental source of data for scientific discovery, even more so in a situation of accelerated research work such as the current pandemic. Our curation of molecular interactions related to *Coronaviridae* enables a systematic perspective of this data and greatly increases its interoperability. From our overview analysis of the data available so far, we can highlight how the SARS-CoV-2 data seems to be both biologically relevant, and thus informative for research on the disease, and strongly preliminary, so full consideration for its inherent incompleteness should be given when using it. The IMEx Consortium is expanding the resource as new data becomes available, striving to provide the most accurate possible picture of the *Coronaviridae* interactome.

## Methods

### Data sources

All stats and details shown on this manuscript are derived from the IntAct database release of 2020-04-30. Complex Portal data used for human complexes overlap checks was also obtained from the same data release.

Specific analyses include the following datasets: SARS-CoV-2 - all human targets of SARS-CoV-2 viral proteins; SARS-CoV-1 - all human targets of SARS-CoV-1 viral proteins; Gordon_LT - all human targets of SARS-CoV-2 viral proteins derived by Gordon et al. plus selected SARS-CoV-2 low throughput studies; Li_LT - all human targets of SARS-CoV-2 viral proteins derived by Li et al. plus selected SARS-CoV-2 low throughput studies. The selected low-throughput studies added to Gordon and Li studies represent just two representative interactions that were not detected in either study, ACE2-Spike and BSG-Spike, so they were not analysed separately. Formatted subsets are available in supplementary table 5.

### Analysis software and packages

Analyses were done using R 4.0.0 (22) and the R data.table package (23). Venn Diagrams were created using ggVennDiagram 0.3 (24). All other plots were created using ggplot23.3.0 (24) and wesanderson (25) packages.

### Tissue enrichment analysis

Analysis was performed using TissueEnrich (19) 1.8.0 R package on human protein targets using Protein Atlas (26) expression data. Background was the entire proteome as listed in the Human Protein Atlas website (www.proteinatlas.org). Only hits with a fold change above zero are shown. Log10 p-value was zero for all tissues. Full data is available in supplementary table 3.

### Pathway enrichment analysis

Pathway enrichment analysis was performed using pathDIP 4 (27) API in R, with literature curated set. Only pathways with false discovery rate <0.01 (BH-method) were considered. As the majority of enriched pathways (75% for SARS-CoV-1, 80% for SARS-CoV-2, 72% for Gordon_LT and 76% for Li_LT) were from Reactome database (28), we further organized them using level one Reactome ontology to create the figure. All enriched pathways are listed in the Supplementary Table 2, and pathways present in multiple sets are highlighted in the “overlap” tab.

## Acknowledgements

The authors would like to thank Dr Yun Niu for assistance in pathway reduction analysis and Dr Serene Wong for useful discussion. The IntAct and Complex Portal databases and EMBL-EBI-based authors received funding from EMBL core funding, Open Targets (grant agreements OTAR-044 and OTAR02-048) and the Wellcome Trust grant INVAR (grant ref: 212925/Z/18/Z). IID and some computational analyses were in part supported by the grants from Ontario Research Fund (#34876), Natural Sciences Research Council (NSERC#203475), Canada Foundation for Innovation (CFI #29272, #225404, #33536), Buchan Foundation and IBM. Work by the UniProt database was supported by the National Eye Institute (NEI), National Human Genome Research Institute (NHGRI), National Heart, Lung, and Blood Institute (NHLBI), National Institute on Aging (NIA), National Institute of Allergy and Infectious Diseases (NIAID), National Institute of Diabetes and Digestive and Kidney Diseases (NIDDK), National Institute of General Medical Sciences (NIGMS), National Cancer Institute (NCI) and National Institute of Mental Health (NIMH) of the National Institutes of Health under Award Number [U24HG007822] (the content is solely the responsibility of the authors and does not necessarily represent the official views of the National Institutes of Health). MatrixDB is supported by the Fondation pour la Recherche Medicale (DBI20141231336), and the Institut Français de Bioinformatique (Glycomatrix project, 2015). DIP Database is supported by the National Institute of General Medical Sciences (R01GM123126). MINT is supported by ERC European Research Council Grant N. 322749 (to G.C.), AIRC startup 21815 and AIRC Project IG 2017 [N.20322 to G.C.]

---

**Supplementary Figure 1: Tissue enrichment analysis of SARS-CoV-2 and SARS-CoV-1 human interacting partners in different data subsets.**

Only tissues with detectable fold change are shown. Horizontal line marks a fold change of 5.

