## Supplementary figures and images for "The IMEx Coronavirus interactome: an evolving map of Coronaviridae-Host molecular interactions"

### Supplementary Figure 1

#### Slide 1
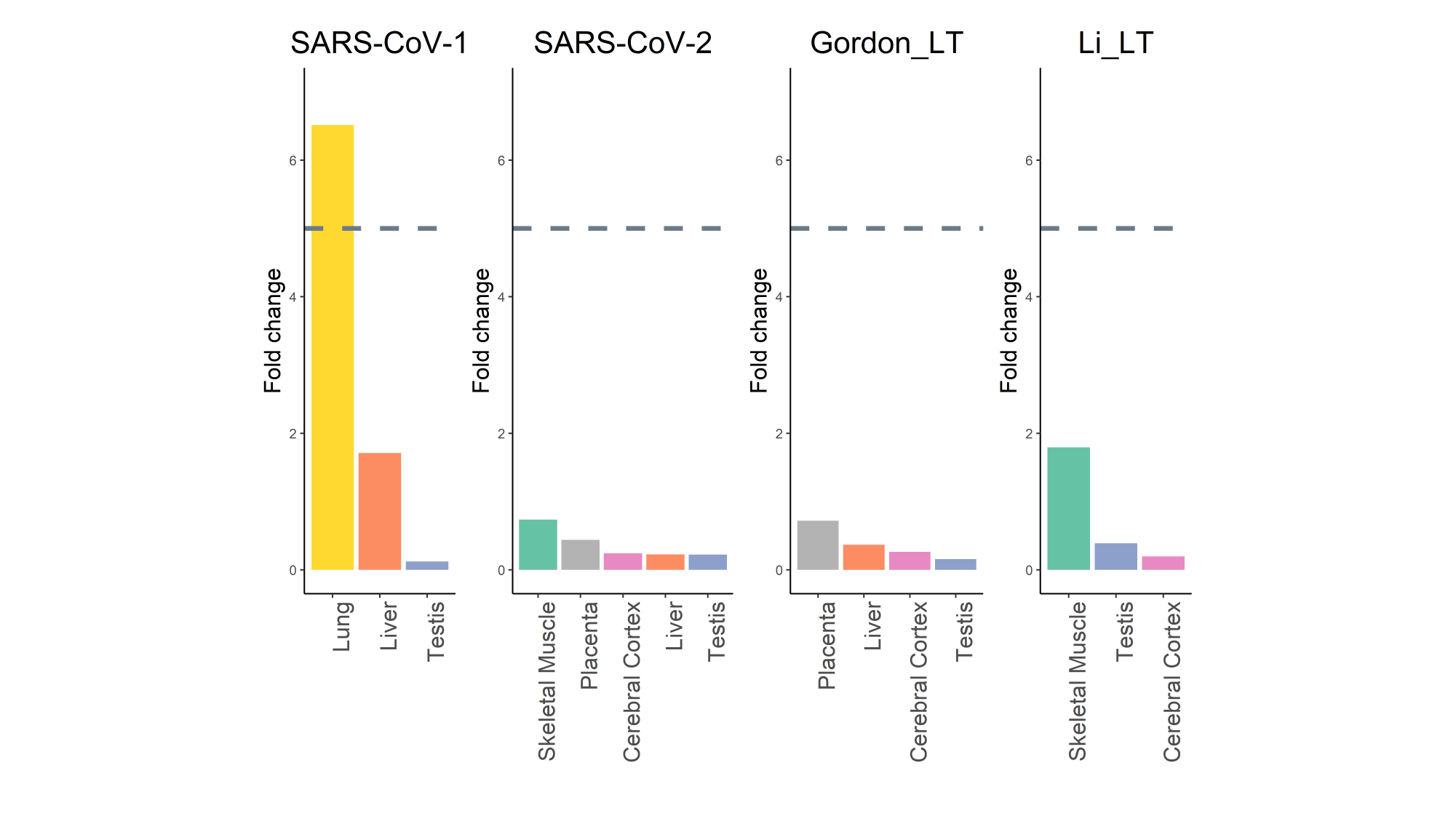
